# Ageing before birth: pace and stability of prenatal growth affect telomere dynamics

**DOI:** 10.1101/809087

**Authors:** Antoine Stier, Neil B. Metcalfe, Pat Monaghan

**Affiliations:** Institute of Biodiversity, Animal Health and Comparative Medicine, University of Glasgow, Glasgow, UK; Department of Biology, University of Turku, Turku, Finland

**Keywords:** Ageing, telomere, embryo, fetal programming, oxidative stress, glucocorticoid

## Abstract

Pre-natal effects on telomere length are increasingly recognized as a potential contributor to the developmental origin of health and adult diseases. While it is becoming clear that telomere length is strongly influenced by pre-natal conditions, the factors affecting telomere dynamics during embryogenesis remain poorly understood. We manipulated both the pace and stability of prenatal growth using incubation temperature in Japanese quail and investigated the impact on telomere dynamics from embryogenesis to adulthood, along with potential drivers of telomere shortening such as oxidative damage and prenatal glucocorticoid levels. Telomere length was not affected by these manipulations for the first 75% of prenatal development, but was reduced at hatching in response to both experimentally induced fast embryo growth and growth instability. These early-life effects on telomere length persisted until adulthood. The effect of developmental instability on telomere length at hatching was potentially mediated by an increased secretion of glucocorticoid hormones during development. Both the pace and the stability of prenatal growth appear as key factors determining telomere length and dynamics up to adulthood, with fast and unstable embryo growth leading to short telomeres with the potential for adverse associated outcomes in terms of reduced longevity and greater risk of disease.

## Introduction

Scientific evidence from both a biomedical (*e.g.* [1,2]) and an ecological perspective (*e.g.* [3,4]) has established that conditions experienced during early life can have profound effects on adult phenotypes and subsequent life histories. For example, accelerated postnatal growth has been shown in a range of species, including humans, to be associated with reduced longevity [5,6] and increased risks of developing numerous age-related pathologies such as cardiovascular disease and type 2 diabetes later in life [2,7]. Telomere shortening (an important hallmark of ageing [8]) has emerged as one of the key candidates linking these early-life conditions to later life adverse effects [9–11]. Telomeres are protective complexes consisting of repeats of a non-coding DNA sequence, proteins and associated RNAs situated at the end of eukaryotic chromosomes. Telomeres serve to identify the chromosome ends, thereby maintaining genome stability, and to prevent the loss of DNA from the chromosome ends that occurs during cell division penetrating into the coding sequences [12]. The length of telomeres is dynamic and results from a balance between loss and restoration processes. In addition to the the ‘end-replication problem’, oxidative stress leading to DNA damage increases telomere loss [13]. While telomere restoration/elongation processes also exist, for instance the enzyme telomerase or the Alternative Lengthening Pathway (ALT), in many species their activity in most somatic tissues is reduced in adulthood most probably as a protection against cancer [14]. Telomeres shorten with age in a broad range of organisms [15,16], with the more pronounced shortening being observed during growth [17,18], presumably because of the intense cellular proliferation and metabolic activity required to sustain growth [11]. Telomere length and/or shortening rate, especially in early life, have been shown to predict subsequent survival/lifespan in a range of species [19–23], thereby leading to the idea that telomere length could act as a biomarker of individual ‘biological state’.

While the main focus of research on telomeres during early life has so far been the postnatal growth period [11], changes during the prenatal period are increasingly recognised as potentially even more important [9,24–28], giving rise to the ‘*fetal programming of telomere biology hypothesis*’ [10]. While the ‘initial’ telomere length (*i.e.* defined by [10] as the telomere length at birth) seems to be of prime importance, the information we have about the prenatal factors determining this ‘initial’ length remain very limited [10].

During the postnatal phase, telomere shortening is thought to be accelerated both by fast growth (*e.g.* [24]) and/or by growing under harsh environmental conditions (*e.g.* [29]), and the same pattern could exist at the prenatal stage. However, most of the experiments to date that have attempted to explore the effect of the pre-natal environment on telomere dynamics have two main limitations. First, measurements of telomere length have either been done at a single time point or using a cross-sectional approach with the first sampling occurring weeks/months after birth (*e.g.* [24–26]), precluding separation of the direct effects during embryogenesis from indirect effects linked to post-natal compensatory responses. Second, experimental manipulation of the pre-natal environment using mammals requires the manipulation to be applied to the mother (*e.g.* [24,26]), leading to potentially confounding effects or compensatory maternal responses (*e.g.* preferential allocation of resources to the embryos).

A good model system for investigating the direct effects of pre-natal conditions on telomere length and dynamics is an oviparous species, where pre-natal conditions can be manipulated directly without interaction with the mother. Telomere length should also be evaluated at different life stages (*i.e.* embryo, birth, during growth and at adulthood) to better understand the impact of prenatal conditions on ‘initial’ telomere length and subsequent telomere shortening rate. In this context, it has been shown that early stage (72hr) embryonic telomere length decreases with ovulation order in captive zebra finches (*Taeniopygia guttata*) and that this effect appears to be maintained until adulthood [27]. Additionally, by incubating eggs of wild common terns (*Sterna hirundo*) at two different temperatures to induce differences in embryo growth rate, Vedder et al. [28] found that slow embryonic growth was associated with slightly longer telomeres at hatching. While the use of stable incubation conditions in that study removed the confounding effect of parental behaviour, constant incubation temperatures are unlikely to be realistic for most egg-laying species [30]. Similarly, variations in the transfer of nutrients, oxygen, hormones from the placenta to the embryos could occur during gestation in mammals and alter embryo development [31]. It is therefore also important to examine the extent to which such unstable developmental conditions could influence embryonic telomere dynamics, and to explore the effects not only on ‘initial’ telomere length, but also on postnatal telomere dynamics. Additionally, it is important to identify the underlying mechanisms driving telomere shortening in response to prenatal conditions. While oxidative stress is a first obvious candidate [13], glucocorticoid hormones have been suggested to influence prenatal telomere shortening [25], in a manner that could be independent of oxidative stress but linked to cellular metabolic reprogramming [32].

The main aim of this study was to examine the effects of prenatal growth rate and developmental stability on telomere dynamics from embryogenesis to adulthood, and to investigate the potential underlying mechanisms (*i.e.* oxidative stress, metabolic rate, prenatal glucocorticoid levels). We used the Japanese quail (*Coturnix japonica*) as a study system for manipulating prenatal growth rate and stability using modulations of incubation temperature, since incubation in this species has been well studied [33], it has a short generation time and is widely used as a model species in physiological, molecular and behavioural studies [34].

## Material and Methods

### Experimental design

All procedures were conducted in accordance with European regulations under the Home Office Project Licence 70/8335 granted to PM and the personal Home Office Licence ICB1D39E7 granted to AS. Japanese quail are precocial birds that can be raised independently from their parents, therefore avoiding any confounding effect linked to parental care. They reach sexual maturity and therefore adulthood very quickly (*ca.* 50-60 days), and are short-lived (2.5-5 years) especially for an avian species of their body size [34]. Japanese quail eggs were obtained from Moonridge Farm (Devon, UK) and delivered within 48 hours of collection. Identity of the parents was unfortunately unavailable, but since eggs were collected every day and Japanese quail lay a maximum of one egg a day [34], it is unlikely that several eggs originated from the same female.

Eggs were incubated in 4 artificial incubators (Brinsea Octagon Advance 20 EX) with automatic egg turning and humidity set at 45% until standardised embryonic day 15 (stED15; see below for description of development times). Egg turning was stopped at stED15 and humidity was increased to 65% for hatching purposes. Experimental temperature conditions were chosen, based on the literature [33] and pilot experiments, so as to maximize differences in developmental speed while minimizing the risks of having differences in hatching success between groups (*i.e.* to avoid the selective disappearance of embryos in some groups). Eggs were incubated at 3 constant temperatures (Fig 1): high (H) = 38.4°C, medium (M) = 37.7°C and low (L) = 37.0°C. Additionally a fourth group was incubated under ‘unstable’ (U) temperature conditions, mimicking the natural incubation pattern, with an incubation temperature of 37.7°C but five incubation recesses of 30min during the day (*i.e.* mimicking the female leaving the nest to forage; [35]). Incubation recesses were achieved by using an automatic power switch; temperature within the incubator during the recesses was measured using a digital thermometer, and averaged over the 5 recesses of one day (averaged temperature is presented in Fig 1, minimum temperature during incubation recess was 29.5°C). While the incubation temperature of this unstable group outside the recess periods was similar to the medium temperature group (37.7°C), its mean daily incubation temperature was 37.0°C, so similar to the low temperature group.

**Fig 1:**
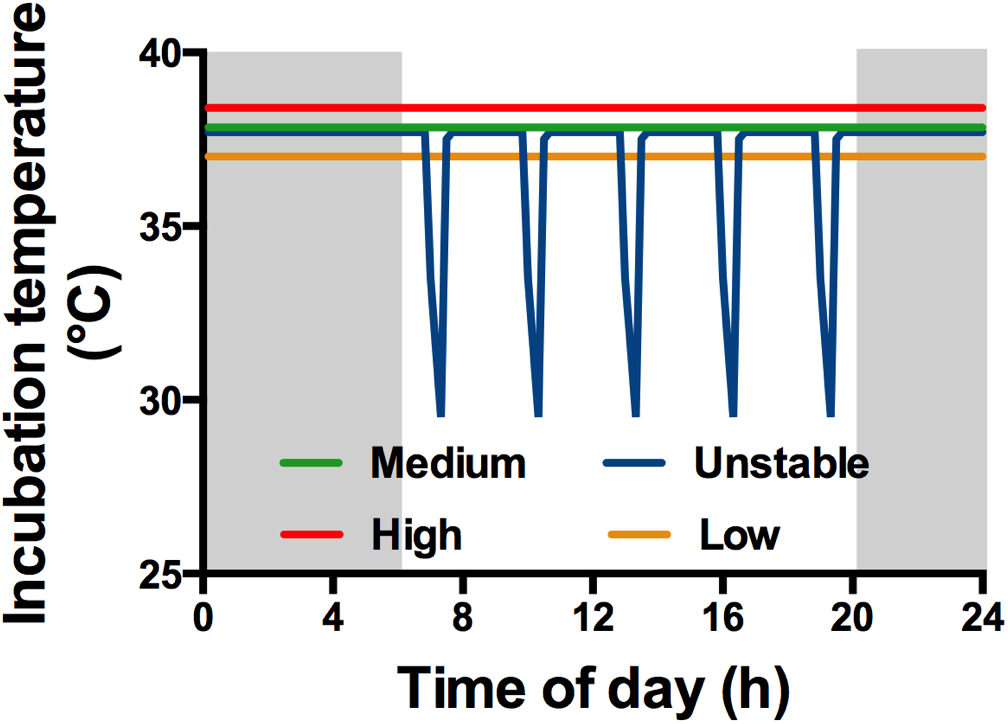
Incubation temperature treatments. Grey and white areas represent dark and light periods respectively. The Unstable group was incubated at the same temperature as the Medium group, but experienced 5 incubation recesses of 30min per day so that its average daily temperature was the same as the Low group.

Animal husbandry rooms were maintained at 21°C on a 14L:10D cycle throughout the experiment, and hatching was monitored during the 14 hours of light starting from stED15 onwards. If hatching occurred overnight, hatching time was considered to be 06:00 am the next morning. Body mass was recorded on the day of hatching (D0) and chicks were placed for 24h in a larger incubator set at 37°C before being placed into their respective enclosure at D1 where an additional heat source was provided until day 15 post-hatching (D15) (Brinsea Comfort brooder 40, 42W), as well as *ad libitum* food (Heygates starter crumbs, 22% protein) and water. We used group-specific enclosures of 4.3m^2^ to avoid late-hatching chicks (from L and U groups) being out-competed by older chicks (from H and M groups) since hatching occurred asynchronously between groups (see results for details). Chick food was switched to pellets (Heygates quail and partridge pellets, 16% protein) at D15. Chicks were maintained in mixed-sex groups until D25 when they could be sexed morphologically. Females were then kept in groups in the enclosures (enclosure size was adjusted to the number of females) and males were placed in pairs in 0.8m^2^ cages to avoid female exhaustion due to male harassment and to limit male-male conflicts. Body mass was recorded for each bird at D0, D1, D3, D5, D10, D15, D20, D30, D45 (by which point birds had reached adult body size), D60 (sexual maturity reached by all birds), D90. Most birds were euthanized at D90-120 for other experimental purposes, but 8 birds were kept alive until approximately one year of age (D360) to gather data on long-term telomere dynamics.

### Sampling procedures

Differences in embryo developmental rate among treatments were initially assessed by measuring embryo mass (after removing yolk and blotting the embryo on absorbent paper) on day 13 of incubation (ED13; 8 eggs sacrificed per treatment). For assessment of embryo telomere length, it was important to collect samples at a similar developmental stage between experimental groups since telomere length is likely to be affected by the number of cell divisions that have occurred (and hence by developmental stage). Since our pilot experiments revealed a difference of approximately 24h in incubation time to hatching between the H and M group, and another 24h between the M and L/U groups, we chose to euthanize and sample embryos at a standardized developmental stage, being the ED13 of the medium temperature treatment (hereafter referred as stED13). Therefore, we euthanized embryos from the high temperature group at ED12 and embryos from the unstable and low temperature groups at ED14 so that all embryos would be approximately at the same developmental stage at the time of sampling. We confirmed the validity of our approach by comparing the body mass of these embryos at stED13, finding no significant differences between groups (H = 4.24 ± 0.13g, M = 4.26 ± 0.16g, L = 4.15 ± 0.13g, U = 4.41 ± 0.19g; N = 14-16 per group; GLM: *F*_3,56_ = 0.5, p = 0.68).

Given that a longitudinal sampling approach is more powerful at revealing changes in telomere length with age than typical cross-sectional sampling, we used red blood cells to measure repeatedly postnatal telomere lengths of the same individuals [16]. We collected blood in embryos at stED13 from the jugular vein using heparinised capillaries following euthanasia by decapitation. Blood was collected at the postnatal stage by puncturing the wing vein with a 26*G* needle and using heparinised capillaries. We collected blood at each of D1, D20 and at D60 for a subsample of 61 of the 103 birds that hatched (+ 8 birds at D360), but blood volume was too low to get enough DNA for TRF telomere analysis for 23 of these 61 chicks at D1. Blood was centrifuged 10min at 3000g and 4°C to separate plasma from RBCs, and was subsequently flash-frozen in liquid nitrogen and stored at −80°C until laboratory analysis. Importantly, telomere length has been shown to be correlated between RBCs and other tissues in birds [36], but a recent study pointed out that it could be life-stage dependent [37]. Therefore, to evaluate the relevance of using RBC telomere length in Japanese quails as a potential indicator of overall telomere length, we compared telomere length in RBCs and in heart samples for a subsample of stED13 embryos (N = 7) and adults (D90; N = 14). Heart samples (10-20mg) were collected from the tip of the heart following euthanasia (cervical dislocation for adults) and dissection on ice, and were quickly flash frozen in liquid nitrogen and stored at −80°C until analysis.

### Embryo heart rate measurement

We measured embryo heart rate as an indicator of embryo metabolism, using a non-invasive infrared methodology (Buddy, Vetronics Services, Devon, UK; [38]) on the 60 eggs being sacrificed at stED13 to control for known effects of developmental stage on embryo heart rate [38]. Heart rate was however measured one day earlier (*i.e.* stED12) to avoid interference with plasma sampling for corticosterone measurement at stED13 (see below). Measurements were always conducted within 90s of taking the egg out of the incubator (*i.e.* to limit cooling down) and never conducted during incubation recess or the following 30min for the Unstable group.

### DNA damage in RBCs

8-OHdG is one of the predominant forms of free radical-induced oxidative lesions in DNA, and widely used as a marker of oxidative damage [39]. Levels of 8-OHdG in RBC DNA was quantified using a competitive immunoassay (300ng DNA, EpiQuick 8-OHdG DNA Damage Quantification Direct Kit Colorimetric, Epigentek, USA) following the manufacturer recommendations. RBC DNA damage is expressed as pg of 8-OHdG/ng of DNA. The intra-individual coefficient of variation based on duplicates was 7.91 ± 1.10% and inter-plate coefficient of variation based on one sample repeated over all plates was 8.38%. A few samples (n = 6) did not contain sufficient DNA for the assay.

### Plasma corticosterone measurement in embryos

Corticosterone (CORT) is the main glucocorticoid hormone in birds. Plasma total CORT levels of embryos at stED13 were determined by immunoassay according to guidelines provided by the manufacturer (DetectX® Corticosterone Enzyme Immunoassay Kit, Arbor Assays, USA). To ensure sampled plasma CORT levels were at baseline levels, the time interval from taking the egg out of the incubator to euthanasia and blood sampling was always kept < 3min (1’40’’ ± 21’’), and embryos from the Unstable group were not sampled during incubation recess or the following 30min. The intra-individual coefficient of variation based on duplicates was 8.00 ± 2.36% and inter-plate coefficient of variation based on one repeated sample over plates was 10.37%.

### Telomere length determination using TRF

Different methods have been used to measure telomere length, the two most common being qPCR and TRF (Terminal Restriction Fragment), the latter being considered as the gold-standard methodology in the field [40]. While trying to validate the qPCR approach for use in Japanese quail by comparing results of qPCR to *in-gel* and denatured TRF (see [41] for details), we noticed the presence of large amounts of interstitial telomeric sequences (ITS), and a pronounced variability in ITS amount between samples. Thereby, methods that could not distinguish between true telomeres and ITS (*i.e.* qPCR and denatured TRF) were found to be poorly indicative of true telomere length in this species, and so we opted for using *in-gel* TRF. We followed a published protocol that has been used successfully in numerous avian species [42]. In brief, DNA was extracted from 5 µL of RBCs using the Gentra Puregene Tissue Kit (Qiagen) and kept frozen at −80°C until analysis. Digestion with restriction enzymes was then conducted on 10µg of DNA using *Hae III* (75U), *Hinf I* (15U), and *Rsa I* (40U) in 1X CutSmart enzyme-buffer and incubated overnight at 37°C. DNA samples and ^32^P labelled size ladder (2-40kb, 1 kb DNA Extension Ladder Invitrogen) were then loaded into a 0.8% non-denaturing agarose gel. DNA was separated using pulse-field gel electrophoresis (14°C at 3 V cm^−1^, initial switch time 0.5 seconds, final switch time 7.0 seconds) for 19 hours, followed by in-gel hybridization overnight with the ^32^P γ-ATP probe (5’-CCCTAA-3’)_4_. Hybridized gels were placed on a phosphor screen for 48 hours, and subsequently scanned with a Typhoon Variable Mode Imager (Amersham Biosciences). Finally, average telomere length was quantified by densitometry in the program ImageJ (version 2.0) within the limits of our molecular size markers (2-40kb). Samples were run over 13 different gels with one standard sample being present in duplicate in each gel to ensure between-gel consistency. Different RBC samples from the same individual (*i.e.* D1, D20 and D60 from a given bird) were always placed on the same gel to ensure the quality of longitudinal data, especially considering the moderate within-individual changes compared to between-individual differences (see ESM1 for details). Experimental groups were balanced within each gel. The intra-gel CV based on duplicates was 2.54 ± 0.40%, and inter-gel CV based on the repeated standard sample was 3.16%.

RBC telomere length decreases with age in Japanese quail from embryogenesis to adulthood (Fig S1A, GEE: Wald-χ^2^ = 177.8, p < 0.001), with the rate of shortening itself decreasing gradually with age (Fig S1B, GEE: Wald-χ^2^ = 87.6, p < 0.001). Based on the 8 individuals sampled twice at adulthood (D60 and D360), the estimated adult telomere shortening rate is −745 ± 109 bp.year^−1^. Japanese quail exhibit a very high intra-individual repeatability/consistency in telomere length (Fig S1C, n = 160, *R adjusted for age effect* = 0.93, 95% CI = [0.90-0.96], p < 0.001). Telomere length in RBCs is significantly correlated with telomere length in the heart (Fig S1D, Spearman σ = 0.80, p < 0.001), a relationship that is found both in embryos (σ = 0.96, p < 0.001) and adult individuals (σ = 0.64, p = 0.015). Telomere length does not significantly differ between heart and RBC samples (exact Wilcoxon test; overall: Z = −1.65, p = 0.10; embryos: Z = −1.69, p = 0.11; adults: Z = −0.63, p = 0.55). Thus telomere length in RBCs can be used as representative of those in other tissues of the quail.

### Data analysis

Data were analyzed using *SPSS* 24 for all statistical tests, except the within-individual repeatability analysis (see above) that was conducted using the *RptR* package in *R* 3.4.2 [43]. Differences in average hatching success between groups were analyzed using a GLMM following a binary distribution. Differences in embryo mass at ED13, embryo heart rate at stED12, incubation time to hatching and plasma CORT were analyzed using GLMs with experimental treatment as a fixed factor, and associated post-hoc tests. We used generalized estimating equations (GEEs) following a normal distribution to investigate the effects of *Age* (*i.e.* repeated effect), *Treatment*, *Sex* and their interactions on body mass dynamics, telomere length dynamics and RBC DNA damage, and the associated post-hoc tests (non-significant interactions were removed from the final models). *Sex* was not included in final models for telomere length and DNA damage since preliminary analyses revealed no significant effect of *Sex* either as a main factor or in interaction with other factors. All means are presented ± SE, and significance level was set at p ≤ 0.05.

## Results

### Developmental and postnatal consequences of incubation temperature and stability

Hatching success did not statistically differ between experimental groups (Fig 2A, Wald-χ^2^ = 1.22, p = 0.75). Experimental treatments altered developmental rate (Fig 2B, GLM: *F*_3,28_ = 49.0, p < 0.001), as shown by the increasing embryo mass at ED13 with increasing incubation temperature (H > M > L, all p < 0.044). Embryos from the unstable group were smaller than the ones from high and medium temperature treatments (all p < 0.001), but not significantly different from the embryos incubated at low temperature (Fig 2B, p = 1.00). Embryo heart rate differed among experimental groups at the same developmental stage (Fig 2C, GLM: *F*_3,56_ = 15.8, p < 0.001), decreasing from high to low temperature (H > M > L, all p < 0.025). Embryos from the unstable group had a heart rate that was intermediate between those from the low and medium temperature groups (Fig 2C, L-U: p = 0.10, M-U: p = 1.0). Incubation time to hatching significantly differed among experimental groups (Fig 2D, GLM: *F*_3,99_= 64.1, p < 0.001). Eggs incubated at lower temperatures hatched later (L > M > H, all p < 0.001), and eggs incubated under unstable conditions hatched later than both high and medium temperatures (all p < 0.001), but were not significantly different in hatching time from low temperature ones (p = 0.92, Fig 2D). However, despite these prenatal differences in developmental rate, there were no overall effects of incubation conditions on postnatal growth (see Table S2, Fig S2 for details on weak age*sex*treatment effects).

**Fig 2:**
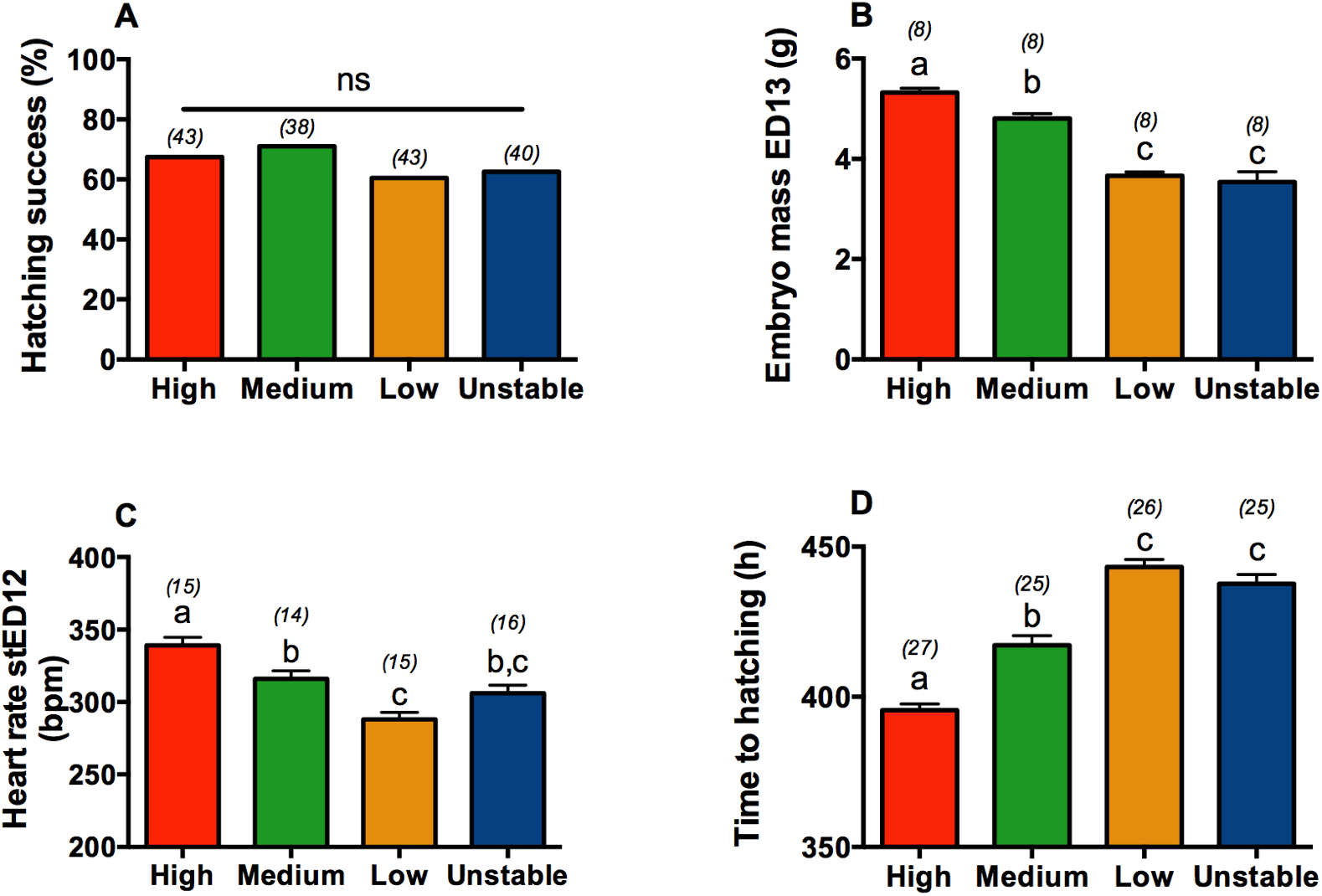
Differences between 4 incubation temperature treatment groups in: (A) Hatching success, (B) Embryo total mass at embryonic day 13 (ED13), (C) Embryo heart rate at standardized embryonic day 12 (stED12), and (D) Time from the beginning of incubation to hatching. Data plotted as means ± SE. Details of statistical tests are given in the text, letters indicate significant differences according to post-hoc tests, and numbers in brackets represent sample size.

### Effect of prenatal conditions on telomere length and dynamics

The experimental manipulation of prenatal conditions significantly influenced telomere length (GEE: Wald-χ^2^ = 8.6, p = 0.034), through an interaction between age and treatment (Fig 3A, GEE: Wald-χ^2^ = 22.0, p = 0.009). Post-hoc tests revealed no significant differences in telomere lengths between groups at the stED13 embryo stage (Fig S3A; all p > 0.45), whereas by day 1 post-hatching both high temperature and unstable chicks exhibited shorter telomeres than did medium and low temperature chicks (Fig S3B; H-M: p = 0.020, H-L: p = 0.020, H-U: p = 0.24, M-L: p = 0.58, M-U: p = 0.003, L-U: p = 0.002). In the middle of the post-natal growth phase (day 20) chicks from the low temperature group still had significantly longer telomeres than unstable and high temperature groups (Fig S3C; L-U: p = 0.015, L-H: p = 0.050), while medium temperature chicks exhibited an intermediate telomere length (Fig S3C; all p > 0.20). At adulthood (day 60), the pattern remained the same as at day 20, with birds from the low temperature group having significantly longer telomeres than unstable and high temperature groups (Fig S3D; L-U: p = 0.007, L-H: p = 0.022), and medium temperature birds having an intermediate telomere length (Fig S3D; all p > 0.13).

**Fig 3:**
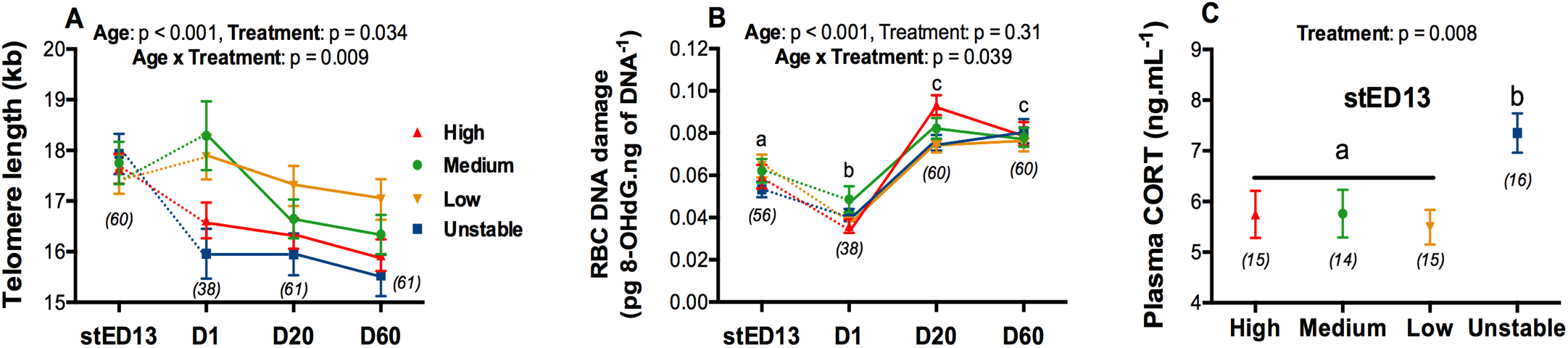
Effects of experimental incubation temperatures on (A) Telomere length and dynamics with age, (B) DNA damage levels and dynamics with age, (C) Plasma corticosterone (CORT) levels in embryos. stED13 represents a standardized developmental stage between experimental groups as explained in the methods; D1 = day after hatching, D20 = chick and D60 = adult. Data plotted as means ± SE. Details of statistical tests are given in the text, solid lines indicate longitudinal data and dotted lines cross-sectional data, letters indicate significant differences according to post-hoc tests, and numbers in brackets represent sample sizes.

The changes in telomere length with age varied significantly between experimental groups (Fig 3A). In both the high temperature (Fig S3E) and unstable (Fig S3H) groups, telomere length decreased during late embryogenesis (from stED13 to day 1 post-hatching; U: p = 0.001; H: p = 0.020) and from juvenile to adulthood (day 20 to 60 post-hatching; U: p < 0.001; H: p < 0.001), but not significantly so during the early chick stage (between day 1 and 20; U: p = 0.95; H: p = 0.32). In contrast, telomere length did not significantly change in the late embryo stage in the medium (Fig S3F) and low (Fig S3G) temperature groups (M: p = 0.48; L: p = 0.55), but subsequently decreased both from day 1 to 20 (M: p = 0.001; L: p = 0.002) and from day 20 to 60 (M: p = 0.003; L: p < 0.001).

### Effect of prenatal conditions on DNA damage and embryo corticosterone levels

The experimental manipulation of prenatal conditions significantly influenced DNA damage through an interaction between age and treatment (Fig 3B, GEE: Wald-χ^2^ = 17.7, p = 0.039). Post-hoc tests revealed no significant differences in DNA damage between groups at the stED13 embryo stage (Fig S4A; all p > 0.098), day 1 post-hatching (Fig S4B, all p > 0.055) or day 60 post-hatching (Fig S4D, all p > 0.30). However, at day 20 post-hatching, DNA damage levels were higher in the high temperature group compared to both low and unstable groups (p < 0.001 and p = 0.002, respectively), while medium temperature chicks exhibited intermediate levels (all p > 0.095, Fig S4C). Age-related variations in DNA damage also slightly differed between experimental groups (Fig S4E-H).

Plasma CORT in embryos significantly differed between experimental groups (GLM: *F*_3,56_ = 4.3, p = 0.008), with embryos from the unstable group having higher CORT levels than other groups (Fig 4, U *vs.* H/M/L: all p < 0.045, differences between H/M/L: all p > 0.96).

## Discussion

Our results highlight the importance of both the pace and stability of prenatal growth in determining the ‘initial’ telomere length (*i.e.* the length at birth), but also in determining how telomere length changes from late embryogenesis to adulthood. Our experimental treatments altered the pace of development and metabolism in the predicted direction without affecting hatching success or postnatal growth rate. Therefore, any effect we observed on telomere lengths is unlikely to be driven by selective mortality or differences in postnatal size/growth pattern between experimental groups. We found no evidence that the impact of prenatal conditions on telomere length was driven by differences in oxidative stress, but increased prenatal exposure to glucocorticoids could be an important factor mediating the impact of developmental instability on telomere length.

### Developmental and postnatal consequences of incubation temperature and stability

The incubation temperature affected the pace of development and metabolism in the expected direction, with higher temperatures increasing embryo metabolic rate (estimated here using heart rate) and growth rate and reducing the time to hatching. Unstable incubation conditions produced a developmental rate that was relatively similar to that of the low incubation temperature group, suggesting that the average daily temperature is the main driver of developmental speed since the U and L groups shared the same daily average. Embryo heart rate in the unstable group was intermediate between that of the low and medium temperature groups, probably because we conducted our measurements outside incubation recess periods (*i.e.* at 37.7°C, the same temperature as the medium group). However, the experimental incubation temperature regimes did not affect either the hatching success or size of the hatched chicks, in contrast to a recent study that found a reduced viability and hatching size of Japanese quail eggs incubated at a lower temperature (36.0°C) than our low temperature treatment (37.0°C) [44].

### Prenatal growth rate and stability as determinants of telomere length and dynamics

While neither embryo growth rate nor the stability of incubation conditions influenced telomere length measured in late embryos (at stED13, *i.e.* 75% of prenatal development), both factors clearly affected perinatal telomere dynamics since embryos growing fast (H group, see [28] for similar results) or under less stable conditions (U group) exhibited shorter telomeres at day 1 post-hatching (*ca.* 5 days later). The fact that telomerase activity declines toward the end of embryo development [45] could be one reason explaining why the effects of our treatments only became apparent by the time of hatching. One initial hypothesis to explain the perinatal shortening observed in response to fast growth (*i.e.* high temperature) or developmental instability (*i.e.* unstable temperature) was that differences in oxidative stress levels might occur between groups, leading to effects on telomere dynamics [13]. Indeed, both high metabolic rate (*i.e.* here linked to high incubation temperature) and variations in metabolic rate resembling the hibernation-arousal situation (*i.e.* here experienced by embryos from the unstable group due to repeated shifts from low to high temperature) have been linked to increased oxidative stress levels [46,47]. However, our results on oxidative damage to DNA (*i.e.* no significant differences between groups in DNA damage before day 20 post-hatching) do not support this hypothesis. Prenatal exposure to glucocorticoids has been shown to accelerate telomere shortening [25], and therefore variation in prenatal glucocorticoid levels linked to incubation temperature stability could contribute to the observed differences in hatching telomere length, since we found that embryos experiencing unstable incubation conditions had higher plasma glucocorticoid levels and shorter telomeres at hatching. This effect on telomeres could be a consequence of either reduced telomerase activity ([48], but see [49]) or reprogramming of cellular metabolism as recently proposed by the *metabolic telomere attrition* hypothesis [32]. However, shorter telomeres at hatching in the high temperature group could not be explained by differences in prenatal glucocorticoid exposure, nor by oxidative stress (see above). Instead, they may be related to the increased rate of cellular division likely associated with their fast metabolism and embryonic growth. Since the size at hatching did not differ between groups, it is possible that fast embryonic growth increased the rate of telomere loss per round of cellular division.

The prenatal treatment also affected early postnatal telomere dynamics, with chicks from the high and unstable temperature treatment groups showing no significant shortening from day 1 to day 20 while low and medium temperature ones did. It is possible that compensatory mechanisms (*e.g.* telomerase activity) were triggered following the initial quick perinatal decline in telomere length observed in these groups, yet without being able to fully compensate for their initially shorter telomeres. Indeed, most of the original differences observed in telomere length at hatching between experimental groups remained relatively unchanged until adulthood, thereby supporting the *fetal programming of telomere biology hypothesis* [10] and proving experimentally the extreme importance of the prenatal stage in determining telomere length over the life course of individuals [10,50].

## Conclusion

Since short telomeres have been associated with reduced chances of survival [19,21] and increased risks of ageing-related diseases [50], our study supports the hypothesis that environmental conditions experienced during embryonic life influence future survival prospects and health state through their effects on ‘initial’ telomere length and subsequent postnatal shortening. We show that both the pace and stability of embryonic growth can affect telomere length, and that the perinatal period appears to be a critical period determining telomere length and dynamics up to adulthood. While the exact mechanisms behind this perinatal telomere shortening and its long-term effects on lifespan and disease risk remain to be identified, our model offers important opportunities to identify these mechanisms and test potential intervention strategies to prevent perinatal telomere shortening.

## Data availability

Dataset used in this manuscript is available at: https://doi.org/10.6084/m9.figshare.9994829.v1

## Author’s contribution

AS designed the study, conducted the experimental work, data analysis and wrote the manuscript. PM and NBM had input on study design and data analysis, and commented on the manuscript.

## Acknowledgements

We are grateful to Mark Haussmann, Winnie Boner and Kate Griffith for the help in setting-up TRF methodology, to Franklin Lo, Becky Shaw and Katie Byrne for their help in collecting body mass data and to Graham Law and his team for taking care of animal husbandry. We are grateful to José Noguera for useful comments on a previous draft. Finally, AS is grateful to the crew of the Marion Dufresnes and the French Polar Institute (IPEV) for hosting him from 56 to 21° South while writing the manuscript. The project was funded by a Marie Skłodowska-Curie Postdoctoral Fellowship (#658085) to AS, and AS was supported by a ‘Turku Collegium for Science and Medicine’ Fellowship at the time of writing.

---

**Fig S1:**
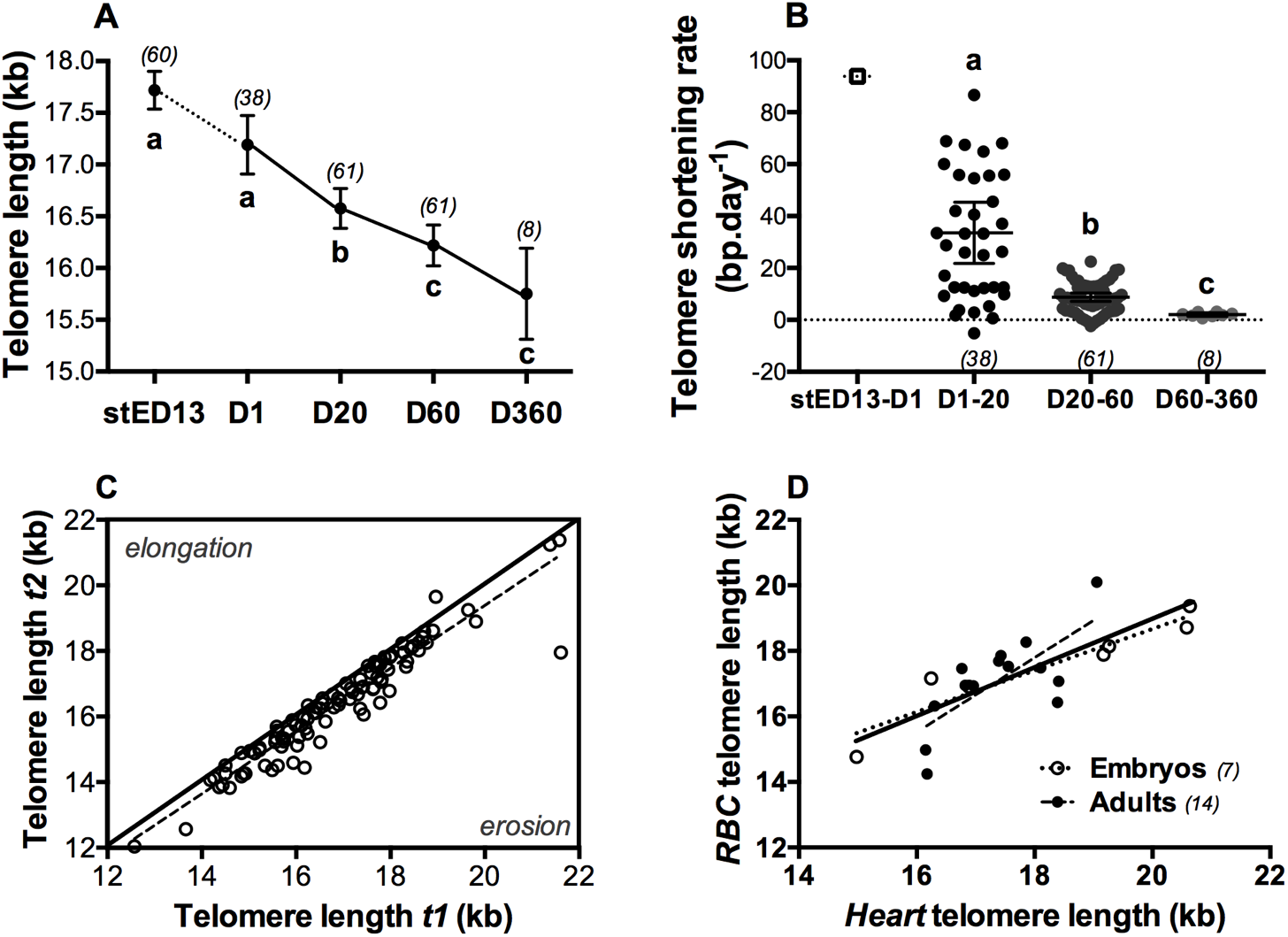
(A) Telomere lengths of Japanese quail red blood cells from embryogenesis to 360 days postnatal. stED13 represents a standardized developmental stage between experimental groups as explained in the main text methods. Data plotted as means ± SE; solid lines linking means indicate longitudinal data and the dotted line cross-sectional data. **(B) Intra-individual telomere shortening rate until 360 days postnatal**. Data plotted as individual data points, with horizontal lines indicating means ± SE. Cross-sectional average ‘shortening rate’ between stED13 and D1 is presented for illustration purposes. **(C) Intra-individual consistency/repeatability of telomere length**. Telomere length at the first sampling occasion (*t1*) is plotted against telomere length at the second sampling occasion (*t2*) for each consecutive measurement (*i.e*. D1 *vs*. D20, D20 *vs*. D60 and D60 *vs*. 360) for each bird. Line of equality is presented as a solid line, while the dotted line represents the relationship in our dataset. **(D) Correlation between heart and RBC telomere length in Japanese quail embryos (stED13) and adult individuals**. Solid line represents the overall correlation while dotted lines represent stage-specific correlations. Details of statistical tests are given in the main text (method section), letters indicate significant differences according to post-hoc tests, and numbers in brackets represent sample sizes.

**Fig S2:**
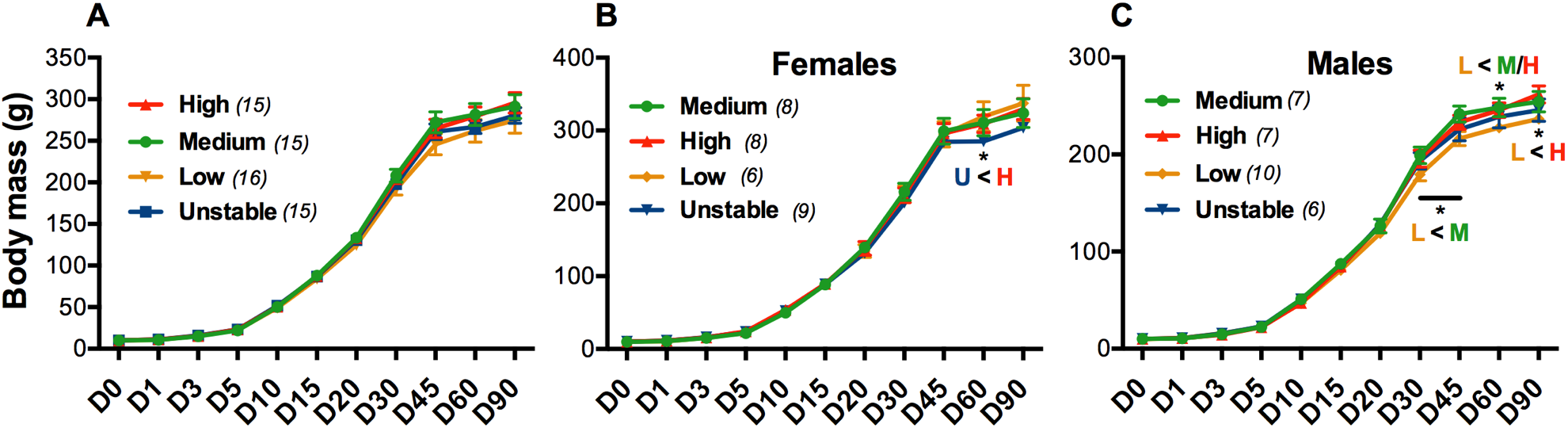
Postnatal body mass growth curves of (A) all individuals, (B) female individuals, (C) male individuals. Data plotted as means ± SE. Details of statistical tests are given in Table S2. * in B and C indicate significant differences according to GEE post-hoc tests. Numbers in brackets represent sample size.

**Fig S3:**
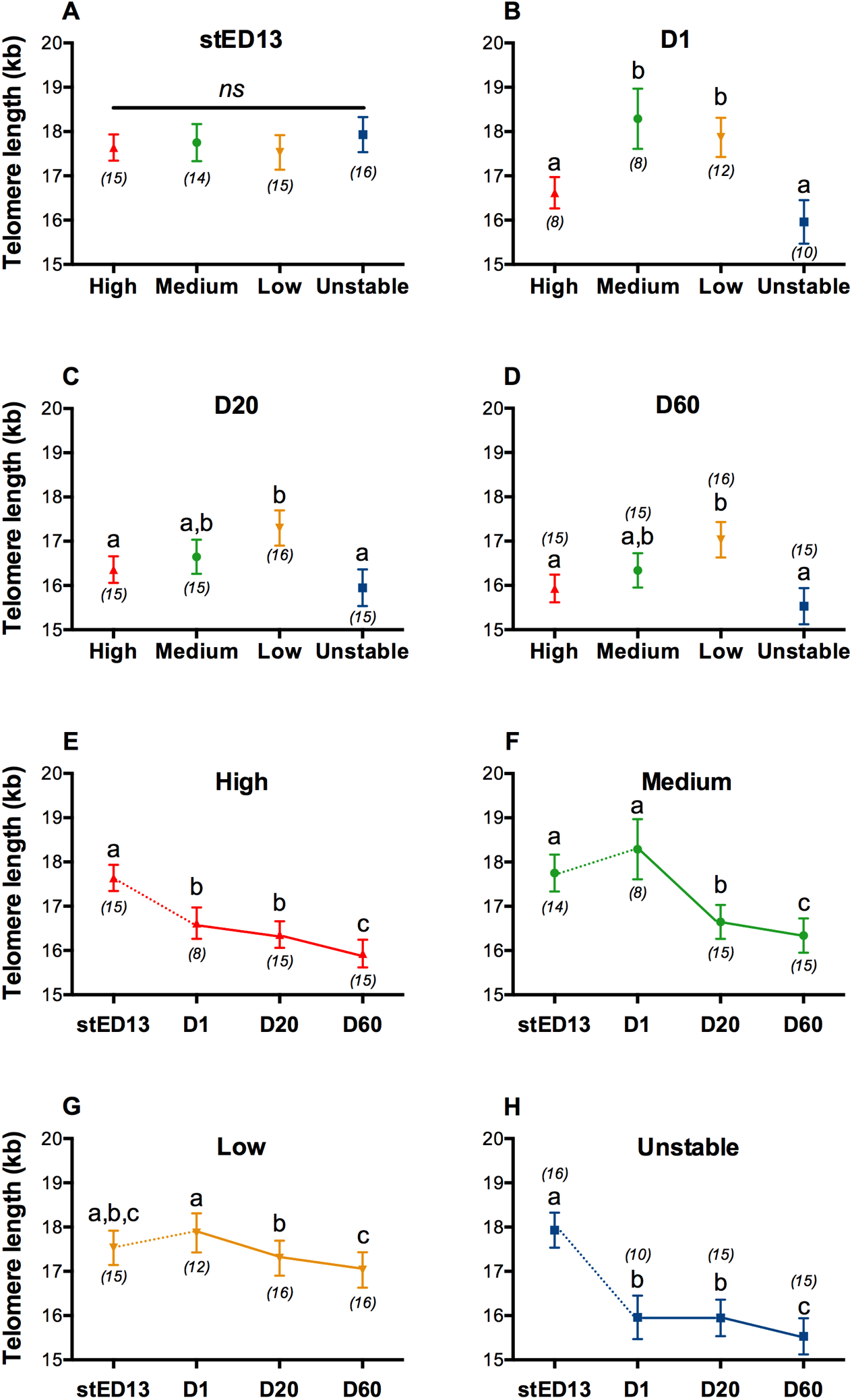
Telomere length in red blood cells according to prenatal experimental conditions and age. stED13 represents a standardized developmental stage between experimental groups as explained in the methods. **(A-D) Telomere length between experimental groups at each stage. (E-H) Age-related changes in telomere length for each experimental group**. Data plotted as means ± SE. Solid lines indicate longitudinal data and dotted lines cross-sectional data, letters indicate significant differences according to post-hoc tests, and numbers in brackets represent sample sizes.

**Fig S4:**
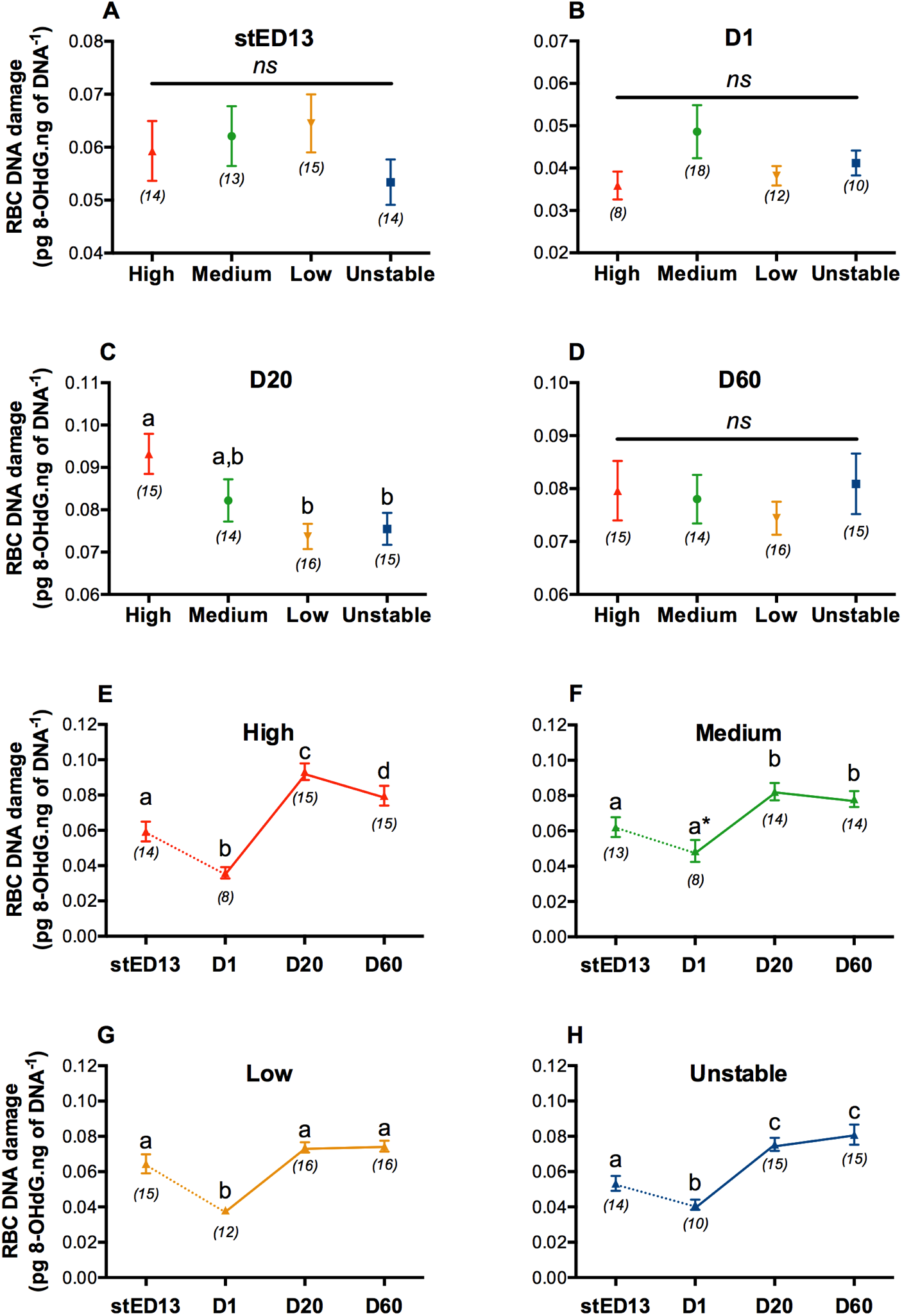
DNA damage levels in red blood cells according to prenatal experimental conditions and age. stED13 represents a standardized developmental stage between experimental groups as explained in the methods. **(A-D) DNA damage levels between experimental groups at each stage. (E-H) Age-related changes in DNA damage for each experimental group**. Data plotted as means ± SE. Solid lines indicate longitudinal data and dotted lines cross-sectional data, letters indicate significant differences according to post-hoc tests, and numbers in brackets represent sample sizes.

**Table S2:** Summary of the most parsimonious GEE models explaining the variability in postnatal body mass growth rate. Estimates for fixed factors are given for the following levels: Age = D0, Treatment = medium and Sex = female. Significant factors are presented in bold.

| **Body mass growth** | **Estimate** | **SE** | **df** | **Wald-$\chi^2$** | **p-value** |
| --- | --- | --- | --- | --- | --- |
| **Intercept** | 245,7 | 10.6 | 1 | 4848.0 | **< 0.001** |
| **Age** | -235.5 | 10.4 | 10 | 6132.3 | **< 0.001** |
| Treatment | 8.6 | 14.4 | 3 | 2.0 | 0.58 |
| **Sex** | 58.1 | 12.3 | 1 | 39.2 | **< 0.001** |
| **Age x Treatment** | -8.6 | 14.1 | 30 | 115.3 | **< 0.001** |
| **Age x Sex** | -58.3 | 12.3 | 10 | 204.7 | **< 0.001** |
| Treatment x Sex |  |  |  |  | ns |
| **Age x Treatment x Sex** | -0.1 | 0.7 | 33 | 63.4 | **< 0.001** |

## References

1. Barker, D. J. & Martyn, C. N. 1991 The maternal and fetal origins of cardiovascular disease. Journal of epidemiology and community health 46, 8–11.

2. Tarry-Adkins, J. L. & Ozanne, S. E. 2017 Nutrition in early life and age-associated diseases. Ageing Research Reviews 39, 96–105. (doi:10.1016/j.arr.2016.08.003)

3. Lindström, J. 1999 Early development and fitness in birds and mammals. Trends in Ecology & Evolution 14, 343–348.

4. Metcalfe, N. & Monaghan, P. 2001 Compensation for a bad start: grow now, pay later? Trends in Ecology & Evolution 16, 254–260.

5. Metcalfe, N. & Monaghan, P. 2003 Growth versus lifespan: perspectives from evolutionary ecology. Experimental Gerontology 38, 935–940.

6. Ozanne, S. E. & Hales, C. N. 2004 Lifespan: Catch-up growth and obesity in male mice. Nature 427, 411–412. (doi:10.1038/427411b)

7. Tarry-Adkins, J. L. & Ozanne, S. E. 2014 The impact of early nutrition on the ageing trajectory. Proc Nutr Soc 73, 289–301. (doi:10.1017/S002966511300387X)

8. López-Otín, C., Blasco, M. A., Partridge, L., Serrano, M. & Kroemer, G. 2013 The hallmarks of aging. Cell 153, 1194–1217. (doi:10.1016/j.cell.2013.05.039)

9. Entringer, S., Epel, E. S., Kumsta, R., Lin, J., Hellhammer, D. H., Blackburn, E. H., Wust, S. & Wadhwa, P. D. 2011 PNAS Plus: Stress exposure in intrauterine life is associated with shorter telomere length in young adulthood. Proceedings of the National Academy of Sciences 108, E513–E518. (doi:10.1073/pnas.1107759108)

10. Entringer, S., de Punder, K., Buss, C. & Wadhwa, P. D. 2018 The fetal programming of telomere biology hypothesis: an update. Philos. Trans. R. Soc. Lond., B, Biol. Sci. 373, 20170151–15. (doi:10.1098/rstb.2017.0151)

11. Monaghan, P. & Ozanne, S. E. 2018 Somatic growth and telomere dynamics in vertebrates: relationships, mechanisms and consequences. Philos. Trans. R. Soc. Lond., B, Biol. Sci. 373, 20160446–15. (doi:10.1098/rstb.2016.0446)

12. De Lange, T., Lundblad, V. & Blackburn, E. H. 2006 Telomeres. Cold Spring Harbor Laboratory Press, New York.

13. Reichert, S. & Stier, A. 2017 Does oxidative stress shorten telomeres in vivo? A review. Biol Letters 13, 20170463–7. (doi:10.1098/rsbl.2017.0463)

14. Shay, J. W., Reddel, R. R. & Wright, W. E. 2012 Cancer and Telomeres-An ALTernative to Telomerase. Science 336, 1388–1390. (doi:10.1126/science.1222394)

15. Gomes, N. M. V., Shay, J. W. & Wright, W. E. 2010 Telomere biology in Metazoa. FEBS Letters 584, 3741–3751. (doi:10.1016/j.febslet.2010.07.031)

16. Stier, A., Reichert, S., Criscuolo, F. & Bize, P. 2015 Red blood cells open promising avenues for longitudinal studies of ageing in laboratory, non-model and wild animals. Experimental Gerontology 71, 118–134. (doi:10.1016/j.exger.2015.09.001)

17. Benetos, A. et al. 2013 Tracking and fixed ranking of leukocyte telomere length across the adult life course. Aging Cell 12, 615–621. (doi:10.1111/acel.12086)

18. Spurgin, L. G., Bebbington, K., Fairfield, E. A., Hammers, M., Komdeur, J., Burke, T., Dugdale, H. L. & Richardson, D. S. 2017 Spatio-temporal variation in lifelong telomere dynamics in a long-term ecological study. Journal of Animal Ecology 282, 20142263–12. (doi:10.1111/1365-2656.12741)

19. Cawthon, R. M., Smith, K. R., O’Brien, E., Sivatchenko, A. & Kerber, R. A. 2003 Association between telomere length in blood and mortality in people aged 60 years or older. The Lancet 361, 393–395.

20. Heidinger, B. J., Blount, J. D., Boner, W., Griffiths, K., Metcalfe, N. B. & Monaghan, P. 2012 Telomere length in early life predicts lifespan. Proceedings of the National Academy of Sciences 109, 1743–1748. (doi:10.1073/pnas.1113306109)

21. Wilbourn, R. V., Moatt, J. P., Froy, H., Walling, C. A., Nussey, D. H. & Boonekamp, J. J. 2018 The relationship between telomere length and mortality risk in non-model vertebrate systems: a meta-analysis. Philos. Trans. R. Soc. Lond., B, Biol. Sci. 373, 20160447–9. (doi:10.1098/rstb.2016.0447)

22. Asghar, M., Hasselquist, D., Hansson, B., Zehtindjiev, P., Westerdahl, H. & Bensch, S. 2015 Chronic infection. Hidden costs of infection: chronic malaria accelerates telomere degradation and senescence in wild birds. Science 347, 436–438. (doi:10.1126/science.1261121)

23. Fairlie, J., Holland, R., Pilkington, J. G., Pemberton, J. M., Harrington, L. & Nussey, D. H. 2015 Lifelong leukocyte telomere dynamics and survival in a free-living mammal. Aging Cell 15, 140–148. (doi:10.1111/acel.12417)

24. Tarry-Adkins, J. L., Martin-Gronert, M. S., Chen, J. H., Cripps, R. L. & Ozanne, S. E. 2008 Maternal diet influences DNA damage, aortic telomere length, oxidative stress, and antioxidant defense capacity in rats. The FASEB Journal 22, 2037–2044. (doi:10.1096/fj.07-099523)

25. Haussmann, M. F., Longenecker, A. S., Marchetto, N. M., Juliano, S. A. & Bowden, R. M. 2012 Embryonic exposure to corticosterone modifies the juvenile stress response, oxidative stress and telomere length. Proceedings of the Royal Society B: Biological Sciences 279, 1447–1456. (doi:10.1098/rspb.2011.1913)

26. Allison, B. J., Tarry-Adkins, J. L., Ozanne, S. E. & Giussani, D. A. 2016 Divergence of mechanistic pathways mediating cardiovascular aging and developmental programming of cardiovascular disease. The FASEB Journal 30, 1968–1975. (doi:10.1096/fj.201500057)

27. Noguera, J. C., Metcalfe, N. B., Reichert, S. & Monaghan, P. 2016 Embryonic and postnatal telomerelength decrease with ovulationorder within clutches. Sci. Rep. 6, 25915. (doi:10.1038/srep25915)

28. Vedder, O., Verhulst, S., Zuidersma, E. & Bouwhuis, S. 2018 Embryonic growth rate affects telomere attrition: an experiment in a wild bird. J Exp Biol 221, 181586. (doi:10.1242/jeb.181586)

29. Stier, A., Delestrade, A., Bize, P., Zahn, S., Criscuolo, F. & Massemin, S. 2016 Investigating how telomere dynamics, growth and life history covary along an elevation gradient in two passerine species. Journal of Avian Biology 47, 134–140. (doi:10.1111/jav.00714)

30. Deeming, D. C. 2002. Nests, eggs, and incubation: new ideas about avian reproduction. Oxford Ornithology Series.

31. Fowden, A. L., Forhead, A. J., Coan, P. M. & Burton, G. J. 2008 The Placenta and Intrauterine Programming. J Neuroendocrinol 20, 439–450. (doi:10.1111/j.1365-2826.2008.01663.x)

32. Casagrande, S. & Hau, M. 2019 Telomere attrition: metabolic regulation and signalling function? Biol Letters 15, 20180885–11. (doi:10.1098/rsbl.2018.0885)

33. Romao, J. M., Moraes, T. G. V. de, Teixeira, R. S. de C., Buxade, C. C. & Cardoso, W. M. 2010 Incubation of Japanese quail eggs at different temperatures: hatchability, hatch weight, hatch time and embryonic mortality. Archives of Veterinary Science 14, 155–162.

34. Ottinger, M. A. 2001 Quail and other short-lived birds. Experimental Gerontology 36, 859–868.

35. Orcutt, F. S., Jr & Orcutt, A. B. 1976 Nesting and parental behavior in domestic common quail. The Auk 93, 135–141.

36. Reichert, S., Criscuolo, F., Verinaud, E., Zahn, S. & Massemin, S. 2013 Telomere Length Correlations among Somatic Tissues in Adult Zebra Finches. PLoS ONE 8, e81496. (doi:10.1371/journal.pone.0081496.t004)

37. Schmidt, J. E., Sirman, A. E., Kittilson, J. D., Clark, M. E., Reed, W. L. & Heidinger, B. J. 2016 Telomere correlations during early life in a long-lived seabird. Experimental Gerontology 85, 28–32. (doi:10.1016/j.exger.2016.09.011)

38. Sheldon, E. L., McCowan, L. S. C., McDiarmid, C. S. & Griffith, S. C. 2018 Measuring the embryonic heart rate of wild birds: An opportunity to take the pulse on early development. The Auk 135, 71–82. (doi:10.1642/AUK-17-111.1)

39. Halliwell, B. & Gutteridge, J. 2007 Free Radicals in Biology and Medicine. Oxford: Oxford University Press.

40. Nussey, D. H. et al. 2014 Measuring telomere length and telomere dynamics in evolutionary biology and ecology. Methods in Ecology and Evolution 5, 299–310. (doi:10.1016/S0968-0004(02)02110-2)

41. Foote, C. G., Vleck, D. & Vleck, C. M. 2013 Extent and variability of interstitial telomeric sequences and their effects on estimates of telomere length. Mol Ecol Resour 13, 417–428. (doi:10.1111/1755-0998.12079)

42. Tricola, G. M. et al. 2018 The rate of telomere loss is related to maximum lifespan in birds. Philos. Trans. R. Soc. Lond., B, Biol. Sci. 373, 20160445–11. (doi:10.1098/rstb.2016.0445)

43. Stoffel, M. A., Nakagawa, S. & Schielzeth, H. 2017 rptR: repeatability estimation and variance decomposition by generalized linear mixed-effects models. Methods in Ecology and Evolution 8, 1639–1644. (doi:10.1111/2041-210X.12797)

44. Ben-Ezra, N. & Burness, G. 2016 Constant and Cycling Incubation Temperatures Have Long-Term Effects on the Morphology and Metabolic Rate of Japanese Quail. Physiol Biochem Zool 90, 96–105. (doi:10.5061/dyad.76qt1)

45. Ulaner, G. A., Hu, J.-F., Vu, T. H., Giudice, L. C. & Hoffman, A. R. 1998 Telomerase Activity in Human Development Is Regulated by Human Telomerase Reverse Transcriptase (hTERT) Transcription and by Alternate Splicing of hTERT Transcripts. Cancer Res 58, 4168–4172.

46. Hermes-Lima, M. & Zenteno-Savín, T. 2002 Animal response to drastic changes in oxygen availability and physiological oxidative stress. Comparative Biochemistry and Physiology Part C 133, 537–556.

47. Stier, A., Bize, P., Habold, C., Bouillaud, F., Massemin, S. & Criscuolo, F. 2014 Mitochondrial uncoupling prevents cold-induced oxidative stress: a case study using UCP1 knockout mice. J Exp Biol 217, 624–630. (doi:10.1242/jeb.092700)

48. Choi, J., Fauce, S. R. & Effros, R. B. 2008 Reduced telomerase activity in human T lymphocytes exposed to cortisol. Brain, Behavior, and Immunity 22, 600–605. (doi:10.1016/j.bbi.2007.12.004)

49. Beery, A. K., Lin, J., Biddle, J. S., Francis, D. D., Blackburn, E. H. & Epel, E. S. 2012 Chronic stress elevates telomerase activity in rats. Biol Letters 8, 1063–1066. (doi:10.1098/rsbl.2012.0747)

50. Aviv, A. & Shay, J. W. 2018 Reflections on telomere dynamics and ageing-related diseases in humans. Philos. Trans. R. Soc. Lond., B, Biol. Sci. 373, 20160436–8. (doi:10.1098/rstb.2016.0436)

